# Mechanism of innate immune reprogramming by a fungal meningitis pathogen

**DOI:** 10.1101/2021.09.02.458767

**Authors:** Eric V. Dang, Susan Lei, Atanas Radkov, Hiten D. Madhani

## Abstract

How deadly fungal pathogens overcome mammalian innate immunity is largely unknown. *Cryptococcus neoformans*, the most common cause of fungal meningitis, induces a pathogenic type 2 response characterized by pulmonary eosinophilia and alternatively activated macrophages. Using forward genetics, we identified a fungal secreted protein, Cpl1, necessary and sufficient to enhance alternative activation of primary macrophages *in vitro*. Cpl1-enhanced polarization requires Toll-like receptor 4, a known mediator of allergen-induced type 2 responses. Cpl1 is essential for virulence, drives polarization of interstitial macrophages *in vivo*, and requires type 2 cytokine signaling for its impact on infectivity. *C. neoformans* selectively associates with polarized interstitial macrophages during infection, supporting a direct hostpathogen interaction. This work identifies a secreted effector produced by a human fungal pathogen that reprograms innate immunity to enable tissue infection.

**One sentence summary:** Identification of a secreted fungal effector that promotes virulence by enhancing type 2 inflammation

## Article

Invasive fungal pathogens are responsible for approximately 1.5 million deaths per year (*1*). They account for 50% of AIDS-related deaths and have been referred to as a ‘neglected epidemic (*2*). New, drug-resistant pathogens such as *Candida auris* have been identified (*3*), while available drugs display limited efficacy and unacceptably high toxicity (*4*). Despite these clinical challenges, little is known about how fungal pathogens evade host immunity to replicate and cause disease. Fungal pathogens of plants utilize secreted effector proteins to thwart plant immune systems (*5*). While secretion of immunomodulatory effector proteins could be a similarly effective strategy for fungi to drive infection in mammals, such molecules have yet to be identified. *Cryptococcus neoformans* is an environmental yeast that is acquired by inhalation and subsequently causes lethal meningitis in immunocompromised individuals (*6*). The most common cause of fungal meningitis, Cryptococcal infection yields case fatality rates that range from 10-70% leading to ~200,000 deaths annually (*7*). In murine infection models, *C. neoformans* induces a type 2 immune response that is detrimental to host protection(*8–12*). Despite the recognized importance of skewed immune responses on the outcome of pulmonary infection, little is known about how *C. neoformans* promotes type 2 inflammation.

Production of a polysaccharide capsule helps *C. neoformans* evade macrophage phagocytosis(*13*). Capsular polysaccharides may have additional immunomodulatory functions (*14*). To test whether capsule contributes to suppression of classical macrophage activation, we measured TNF protein levels in the supernatants of murine bone marrow-derived macrophages (BMDMs) exposed to wild type or capsule-deficient (*cap60Δ*) *C. neoformans* (KN99α serotype A strain), the non-pathogen *Saccharomyces cerevisiae* (s288c), or the pathogen *Candida albicans* (SC5314). *C. neoformans* failed to induce TNF secretion, whereas *S. cerevisiae* and *C. albicans* induced robust production of this cytokine (**Fig. 1A**). To investigate BMDM responses globally, we performed RNA-seq on cells stimulated with LPS, zymosan (a product of *S. cerevisiae* cell walls), *S. cerevisiae*, and wild type or capsule-deficient *C. neoformans*. Consistent with our ELISA data, *C. neoformans* induced negligible induction of inflammatory cytokine mRNAs (**Supp Fig. 1A**). Analysis of genes upregulated by *C. neoformans* revealed a partial signature reminiscent of alternatively activated/tolerized macrophages (sometimes referred to as M2 polarized macrophages), especially the striking induction of the key marker gene arginase-1 (*Arg1*), by both wild type and capsule-deficient strains. (**Fig. 1B and Supp Fig. 1B**). There was minimal induction of *Arg1* by LPS, zymosan, or *S. cerevisiae* treatment (**Fig. 1C and Supp Fig. 1B**). These results suggested that *C. neoformans* harbors a capsule-independent immunomodulatory mechanism.

**Figure 1.**
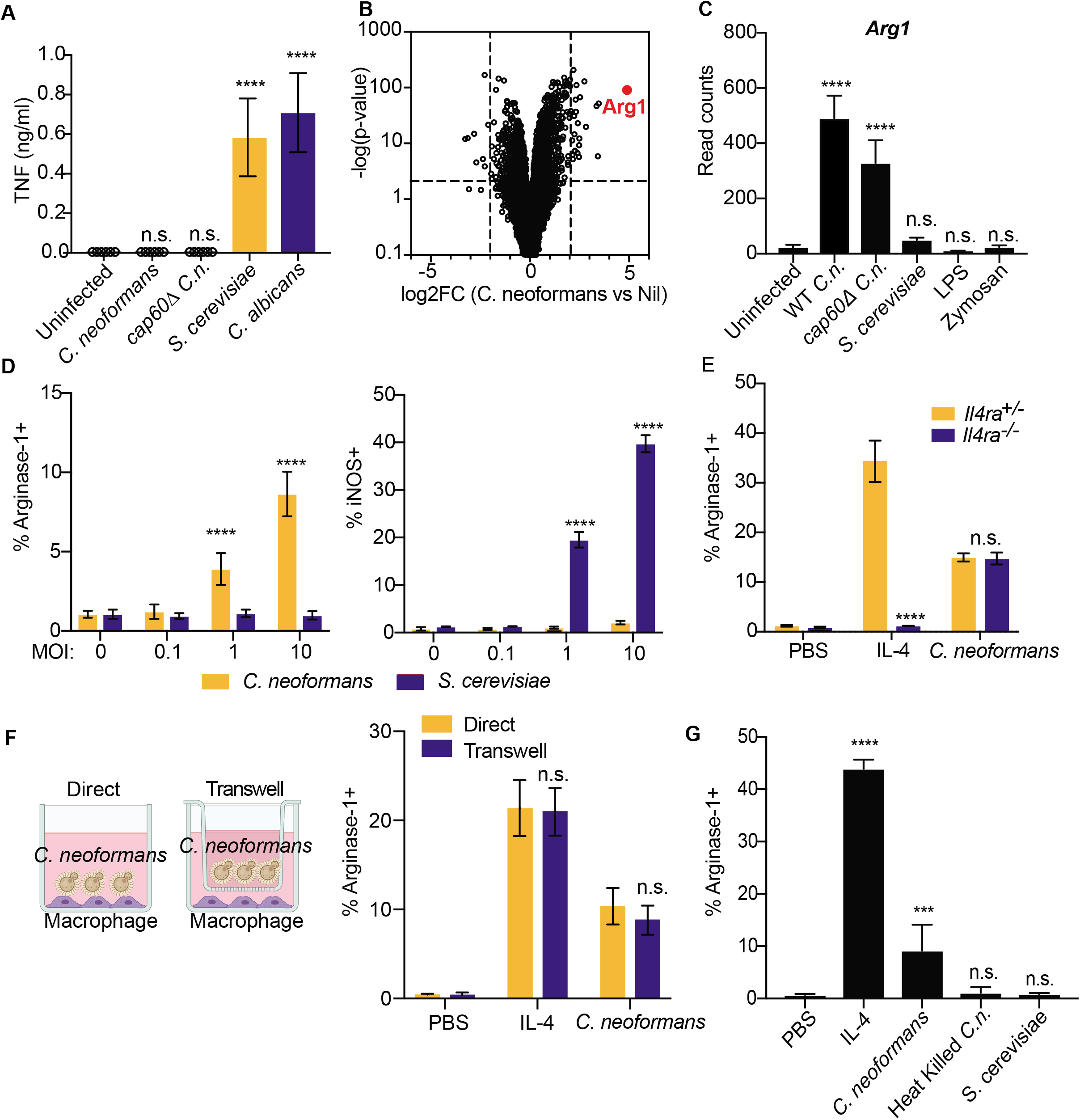
*Cryptococcus* promotes arginase-1 expression in macrophages via a soluble, capsule-independent mechanism. **(A)** TNF ELISA on supernatants from BMDMs infected for 24hrs with the indicated yeasts at MOI=10. **(B)** Volcano plot of RNA-seq data showing differentially expressed genes between uninfected and 6hrs wild type *C. neoformans* MOI=10 infected BMDMs based on DESeq2 analysis. **(C)** Normalized DEseq2 RNA-seq read counts on STAR aligned transcripts from RNA-seq on BMDMs stimulated for 6hrs with either WT *C.n., cap60Δ C.n., S. cerevisiae* (all at MOI=10), LPS (100ng/ml), or zymosan (10ug/ml). **(D)** Intracellular FACS staining for arginase-1 (left) and iNOS (right) after 24hrs of infection with either *C. neoformans* or *S. cerevisiae* at the indicated MOIs. **(E)** Arginase-1 intracellular FACS on either *Il4ra*^+/-^ or *Il4ra*^-/-^ BMDMs stimulated for 24hrs with IL-4 (40ng/ml) or WT *C.n*. (MOI=10). **(F)** Arginase-1 intracellular FACS with identical conditions as in **(E)**, but with the stimuli either added directly to the BMDMs or added to the top of a 0.2um transwell insert (image created with BioRender.com). **(G)** Arginase-1 FACS on BMDMs stimulated for 24hrs with IL-4 (40ng/ml), live WT *C.n*., heat killed (55C for 15min) WT *C.n*., or *S. cerevisiae* (all MOI=10). ***p < 0.001; ****p < 0.0001 by one-way ANOVA with Bonferroni test.

To verify this result, we performed intracellular flow cytometry staining for arginase-1 and iNOS protein and confirmed specific upregulation by IL-4 and LPS, respectively (**Supp Fig. 1C**). *C. neoformans* induced arginase-1 protein in BMDMs in an MOI-dependent manner, whereas no induction was observed with *S. cerevisiae* (**Fig. 1D**). Reciprocally, *Saccharomyces* drove high levels of iNOS, whereas minimal induction was seen after *C. neoformans* infection (**Fig. 1D**). Type 2 cytokines such as IL-4 and IL-13 are well-known to promote expression of Arg1 in macrophages, thus we tested whether *C. neoformans-mediated* Arg1 expression required IL-4Rα. However, *C. neoformans* induced similar levels of arginase-1 in wild type and *Il4ra*^-/-^ BMDMs, suggesting an alternative mechanism (**Fig. 1E**). We found that the *C. neoformans-* derived induction signal was soluble and did not require direct cell-cell interactions, as the fungi could still promote arginase-1 expression across a 0.2um transwell (**Fig. 1F**). However, heat-killed *C. neoformans* could not induce macrophage arginase-1, indicating an active process in live fungi is required for their ability to reprogram the macrophage inflammatory state (**Fig. 1G**).

To obtain a more detailed molecular understanding of how *C. neoformans* influences macrophage polarization, we performed a forward genetic screen using a *C. neoformans* knockout collection generated in the KN99α strain background by our laboratory. To generate this collection, most non-essential genes (n=4,402) were individually replaced with a nourseothricin-resistance cassette (*NAT1*) by homologous recombination. We infected BMDMs with this collection in an arrayed 96-well format and then screened them by flow cytometry to identify *C. neoformans* mutant strains that lost the ability to induce macrophage arginase-1 expression (**Fig. 2A**). We ranked each mutant by Z-score, re-screened the top 100 hits (**Supp Fig. 2A**), and identified 14 genes whose deletion yielded a reproducible defect in arginase-1 induction in response to *C. neoformans* (**Supp Fig. 2B**). Because we had already observed that the *C. neoformans*-derived signal was soluble, we focused on the one protein among the 14 that harbored a predicted signal peptide, which is encoded by *CNAG_02797/CPL1*. CPL1 is a predicted small, secreted protein of unknown molecular function with a cysteine-rich C-terminal domain (**Fig. 2B**). CPL1 was previously identified in an *in vivo* mouse screen of a small knockout library in a different yeast strain background, and was found to have defect in fitness as well as a defect in the production of visible polysaccharide capsule (*15*). As effector proteins from plant fungal pathogens are highly enriched for small, secreted cysteine-rich proteins, we selected CPL1 for further study(*16*).

**Figure 2.**
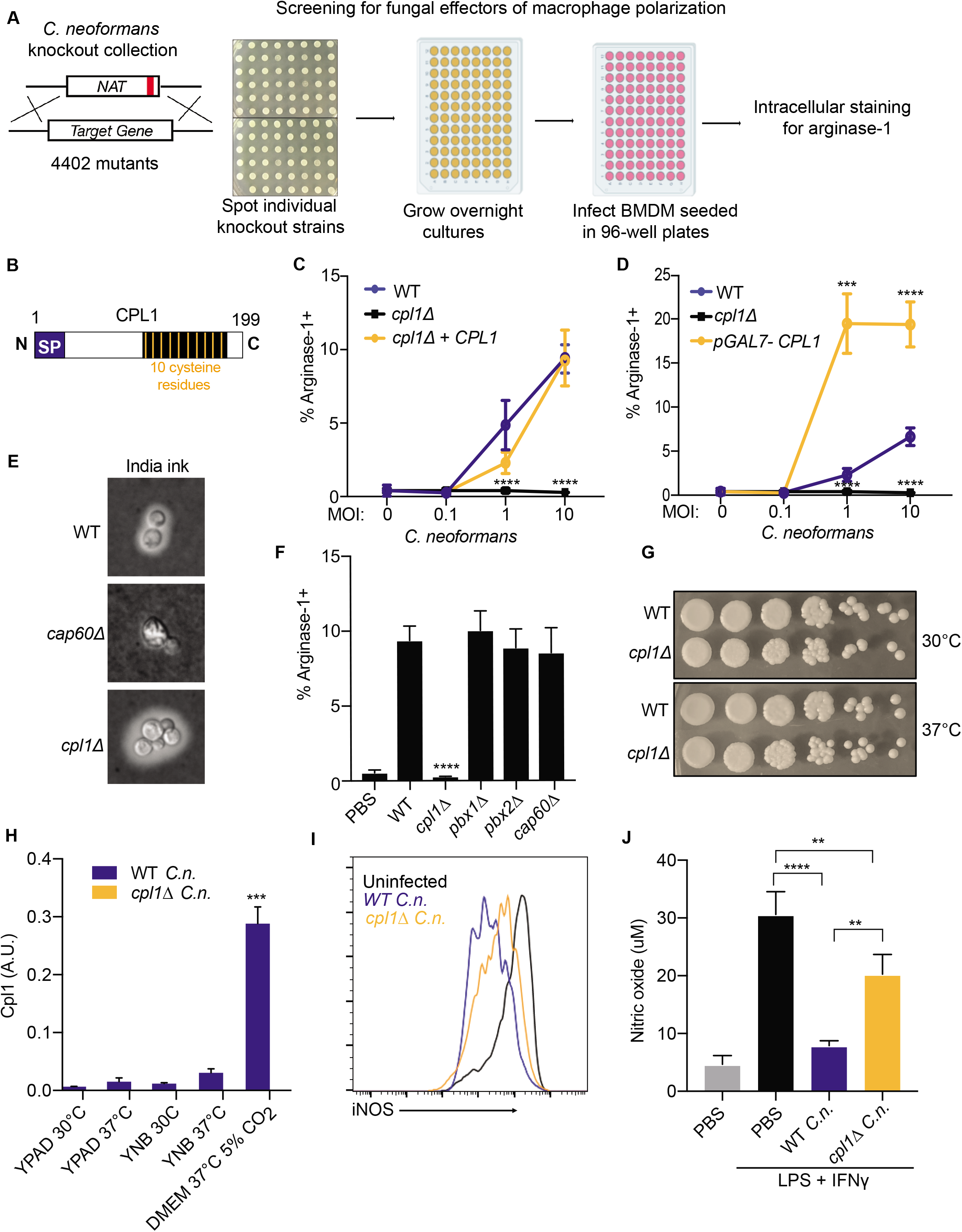
Identification of CPL1 as a fungal effector by forward genetics. **(A)** Schematic of FACS-based forward genetic screening strategy to identify fungal effectors that drive arginase-1 expression. **(B)** Outline of CPL1 protein domain architecture. **(C)** Complementation assay using arginase-1 FACS on BMDMs stimulated for 24hrs with either WT, *cpl1Δ*, or *cpl1Δ* + CPL1 *C.n*. strains at the indicated MOIs. **(D)** Arginase-1 FACS on BMDMs stimulated for 24hrs with WT, *cpl1Δ*, or pGAL7-CPL1 *C.n*. strains at the indicated MOIs. **(E)** Brightfield images of India ink staining for capsular polysaccharides in the indicated strains after overnight culture in 10% Sabouraud media **(F)** Arginase-1 FACS on BMDMs stimulated for 24hrs with the indicated capsule mutant strains at MOI=10. **(G)** Spotting assay for WT vs *cpl1Δ C.n*. growth on YPAD plates incubated at the indicated temperatures. (H) RT-qPCR for CPL1 mRNA expression in cultures grown to OD_600_=1.0 in the indicated conditions (A.U. = arbitrary units relative to ACT1). **(I)** Representative FACS histogram for intracellular iNOS protein levels in BMDMs pre-infected with the indicated strains at an MOI=10 for 2hrs followed by 24hr stimulation with LPS (100ng/ml) and IFNγ (50ng/ml). **(J)** Total nitric oxide in supernatants from BMDMs treated as in **(I)**. **p < 0.01; ***p < 0.001; ****p < 0.0001 by one-way ANOVA with Bonferroni test.

To complement the *cpl1*Δ phenotype, we inserted the *CPL1* gene into a neutral locus (*17*). This complemented strain displayed restored arginase-1 induction capacity (**Fig. 2C**). In addition, overexpression of *CPL1* triggered increased arginase-1 induction (**Fig. 2D**). Experiments performed in different *C. neoformans* strain backgrounds have yielded different conclusions regarding the generality of the capsule defect of *cpl1*Δ mutants (*18, 19*). We found that, in the KN99α background used here, both India ink staining and FACS staining of yeast with antibodies against GXM, the major capsular polysaccharide, showed that the *cpl1Δ* strain contains equivalent to slightly increased surface GXM levels compared to wild type, whereas *cap60Δ* showed an expected loss of staining (**Fig. 2E and Supp Fig. 2C**). While we did not expect capsule production to be related to the arginase-1 induction phenotype, as *cap60Δ* showed equivalent upregulation to wild type by RNA-seq and no classical capsule mutants were obtained in our screen, we nonetheless tested a series of capsule-related mutants that clustered with *cpl1Δ* in chemogenomic profiles (*18*). None showed altered arginase-1 upregulation by BMDMs (**Fig. 2F**). Thus, the phenotype of *cpl1*Δ mutants on arginase-1 induction could not be explained by a defect in the capsule.

In addition to producing a capsule, *C. neoformans* has virulence attributes including the production of melanin, which is thought to have antioxidant properties, and the ability to grow at 37°C (*20*). To determine whether *CPL1* had an effect on growth at mammalian body temperature, we performed spotting assays on YPAD plates grown at either 30°C or 37°C. We observed no specific defect for *cpl1*Δ in growth at 37°C (**Fig. 2G**). We next asked whether temperature or culture conditions had any impact on *CPL1* expression, as genes related to virulence may be predicted to have increased expression in mammalian tissue culture conditions. Indeed, we found that mammalian tissue culture conditions (37°C, DMEM, 10% fetal calf serum, 5% CO_2_) dramatically increased CPL1 mRNA levels compared to culture in YPAD or YNB at either 30°C or 37°C (**Fig. 2H**). We also spotted WT and *cpl1*Δ onto L-DOPA plates to assess melanin production and found qualitatively equivalent melanin induction (**Supp Fig. 2D**).

*C. neoformans* can suppress nitric oxide production by macrophages in response to LPS and IFNγ via a capsule-independent mechanism (*20*). As alternative macrophage activation antagonizes induction of antimicrobial factors such as iNOS, we tested whether *CPL1* had an impact on iNOS expression and nitric oxide production. We infected BMDMs for 2hrs with either wild-type or *cpl1*Δ *C. neoformans* followed by stimulation with LPS and IFNγ for 24hrs. We then used flow cytometry to assess iNOS expression and quantified total nitric oxide in the BMDM supernatants. Consistent with previous reports, we found that wild-type *C. neoformans* suppresses iNOS expression and nitric oxide production (**Fig. 2I and 2J**). Strikingly, the *cpl1*Δ strain is defective in suppressing iNOS expression and nitric oxide production (**Fig. 2I and 2J**). This effect was strong but not complete, indicating that additional modulators exist.

We hypothesized that Cpl1 might act directly on macrophages after secretion by *C. neoformans*. To test this model, we expressed recombinant CPL1-6xHis (rCPL1) using the yeast *Pichia pastoris* (**Fig. 3A**). We then stimulated macrophages with dilutions of purified protein, or the equivalent dilutions derived from mock purifications from wild type yeast to control for potential background contaminants. Using flow cytometry, we found that rCPL1 could induce arginase-1 in BMDMs at nanomolar concentrations (**Fig. 3B**). As *C. neoformans* is well-known to induce type 2 cytokines such as IL-4 during pulmonary infection, we tested whether rCPL1 induction of arginase-1, a well-known IL-4-responsive gene, might reflect an ability to enhance the effects of IL-4. We stimulated BMDMs with IL-4 along with dilutions of rCPL1. We found that rCPL1 potentiated the induction of arginase-1 by IL-4 (**Fig. 3B**). RNA-seq on BMDMs stimulated for 24hrs with either PBS, IL-4 alone, rCPL1 alone, or both (IL-4 + rCPL1) confirmed that rCPL1 increased expression of a large subset of IL-4-induced genes, indicating this is not specific to Arg1 expression (**Fig. 3C and Supp Fig. 3A**). Consistent with our FACS analysis, the RNA-seq results showed striking potentiation of IL-4-induced *Arg1* mRNA expression by rCPL1 (**Supp Fig. 3A**). Another gene that showed strong expression potentiation by rCPL1+IL-4 was *Ccl24*, an eosinophil chemoattractant (**Supp Fig. 3A**). Using a transwell migration assay, we found that splenic eosinophils indeed showed increased chemotaxis towards supernatants derived from BMDMs stimulated with IL-4 + rCPL1 compared to IL-4 alone (**Supp Fig. 3B**). We also found that *C. neoformans* showed enhanced growth in supernatants from BMDMs stimulated with IL-4 + rCPL1 compared to supernatants from either stimulation alone (**Fig. 3D**). These results demonstrated that CPL1 enhances the effect of IL-4 on macrophages, generating conditions that are beneficial to fungal growth *in vitro*.

**Figure 3.**
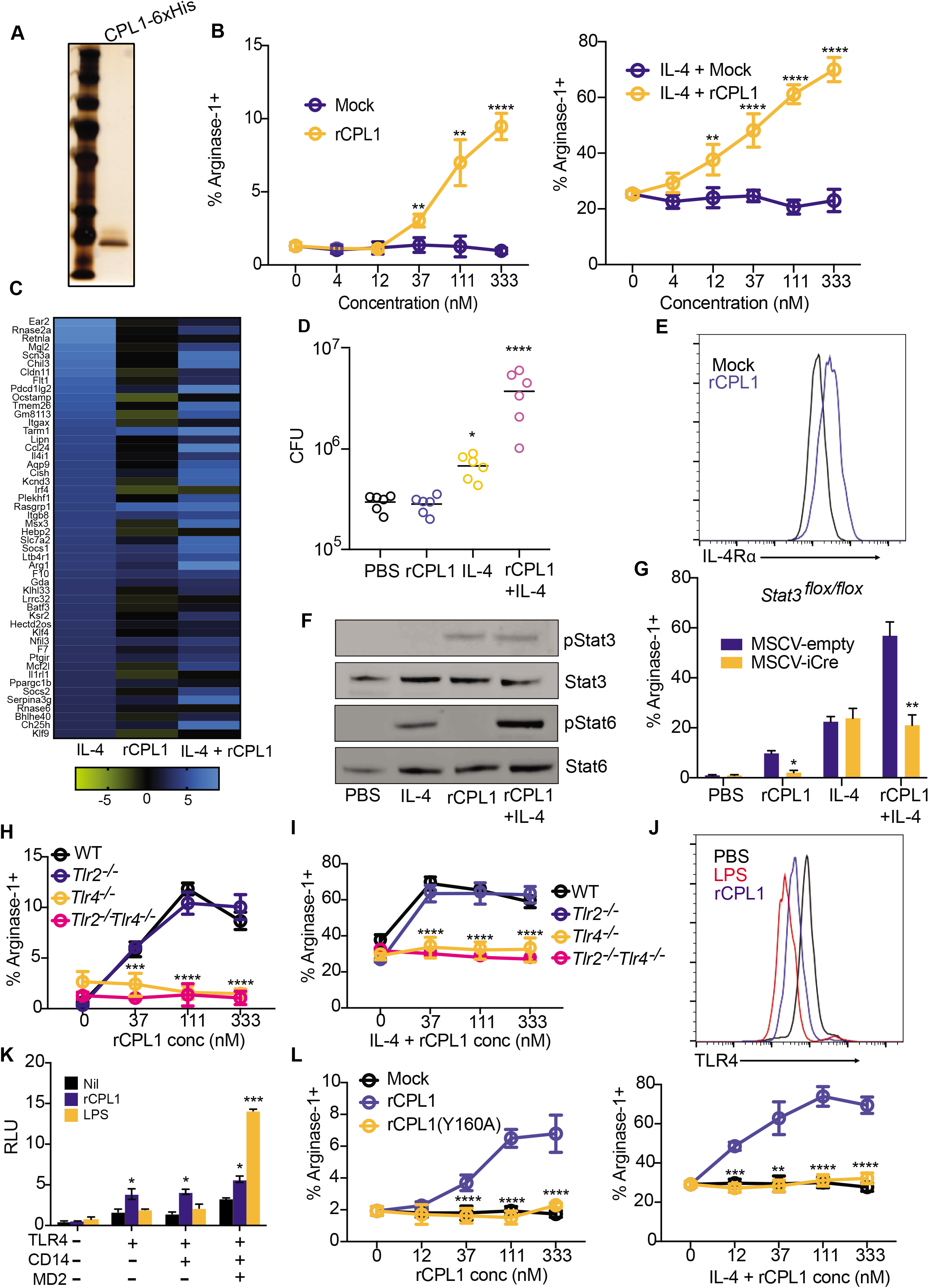
CPL1 activates TLR4/MyD88/STAT3 to potentiate IL-4 signaling. **(A)** Silver stain of SDS-PAGE gel loaded with purified recombinant *P. pastoris* CPL1(6xHis). **(B)** Arginase-1 FACS on BMDMs stimulated for 24hrs with the indicated concentrations of mock purifications or purified rCPL1 alone (left) or in combination with 10ng/ml IL-4 (right). **(C)** Heatmap of the top 50 IL-4-induced genes by RNA-seq comparing the log_2_ fold changes of IL-4 (10ng/ml), rCPL1 (111nM), or IL-4 + rCPL1 stimulated cells to unstimulated cells. **(D)** C. neoformans CFUs after 48hrs of incubation at 37°C, 5% CO_2_ in supernatants from BMDMs stimulated for 24hrs as in **(C)**. **(E)** Representative FACS staining of surface IL-4Rα levels on BMDMs stimulated for 24hrs with either WT or *cpl1Δ Cryptococcus* (MOI=10) (left) or with either 111nM rCPL1 or the equivalent dilution of mock purification (right). **(F)** Western blot for pStat3, Stat3, pStat6, or Stat6 on BMDMs stimulated as in **(C)** for 8hrs. **(G)** Arginase-1 FACS on *STAT3^flox/flox^* BMDMs transduced with MSCV-empty or MSCV-iCre retrovirus and stimulated as in **(C).(H)** Arginase-1 FACS in WT, *Tlr2*^-/-^, *Tlr4*^-/-^, or *Tlr2*^-/-^ *Tlr4*^-/-^ BMDMs stimulated for 24hrs with the indicated concentrations of rCPL1. **(I)** Arginase-1 FACS in WT, *Tlr2*^-/-^, *Tlr4*^-/-^, or *Tlr2*^-/-^*Tlr4*^-/-^ BMDMs stimulated for 24hrs with the indicated concentrations of rCPL1 plus IL-4 (10ng/ml). **(J)** Representative FACS histogram of TLR4 staining on BMDMs stimulated for 1hr with either LPS (100ng/ml) or rCPL1 (111nM). **(K)** Luminescence in HEK293T cells transfected with the indicated plasmids along with a NF-kB firefly luciferase plasmid and then stimulated for 6hrs with either rCPL1 (111nM) or LPS (100ng/ml). **(L)** Arginase-1 FACS in BMDMs stimulated for 24hrs with the indicated concentrations of either mock purification, rCPL1, or rCPL1(Y160A) alone (left) or in combination with 10ng/ml IL-4 (right). *p < 0.05; **p < 0.01; ***p < 0.001; ****p < 0.0001 by one-way ANOVA with Bonferroni test.

We investigated how CPL1 and IL-4 may cooperate to amplify the IL-4 transcriptional signature. One intriguing hypothesis was that CPL1 may enhance macrophage sensitivity to IL-4 at the level of the receptor expression. Indeed, FACS staining revealed that CPL1 drove increased surface levels of BMDM IL-4Rα (**Fig. 3E**). *Salmonella typhimurium* has been reported to drive macrophage M2-like polarization via the effector protein SteE, which activates phosphorylation of STAT3 (*21, 22*). We noted in those reports that the readout for M2 polarization was increased IL-4Rα levels, and that STAT3 was required for IL-4Rα upregulation by *Salmonella*. To test whether similar signaling pathways were required for the activity of CPL1, we performed western blots on extracts of BMDMs to assess phosphorylation of STAT3 and STAT6 after stimulation with either IL-4, rCPL1, or IL-4 + rCPL1. While we saw no effect of CPL1 during immediate early time points (**Supp Fig. 3C**), at 8hrs post-stimulation we found that rCPL1 promoted phosphorylation of STAT3 and drove hyper-phosphorylation of STAT6 when given in combination with IL-4 (**Fig. 3F**). To test a genetic requirement for STAT3 in this circuit, we transduced *Stat3*^flox/flox^ bone marrow using murine retrovirus (MSCV) encoding iCre and generated BMDMs. STAT3 was found to be required for both arginase-1 induction and IL-4 potentiation by CPL1 (**Fig. 3G**).

These data support the conclusion that activation of STAT3 phosphorylation is downstream of CPL1 in promoting alternative macrophage activation. We next sought to understand which molecular pathways were required and, specifically, whether there were macrophage-expressed receptors involved in this process. Previous work had shown that Bacillus Calmette Guerrin (BCG) can thwart macrophage intracellular immunity by promoting expression of arginase-1(*23*) by engaging TLR2-MyD88 signaling to activate STAT3, which drives transcription of arginase-1. We therefore tested *MyD88*^-/-^ BMDMs in our arginase-1 induction and potentiation assays. We found that MyD88-deficiency abrogated arginase-1 upregulation by rCPL1 (**Supp Fig. 4A**). We also observed that rCPL1 instead quenched arginase-1 upregulation by IL-4 in *MyD88*^-/-^ BMDMs, which may be due to induction of type I IFN downstream via the adaptor TRIF (**Supp Fig. 4A**). We hypothesized that MyD88 signaling promoted STAT3 phosphorylation via induction of autocrine/paracrine-acting cytokines. As predicted by such a model, we found that co-culture of CD45.2+ *MyD88*^-/-^ BMDMs with wild type congenic (CD45.1+) BMDMs rescued the arginase-1 induction (**Supp Fig. 4B**).

As MyD88 is an adaptor molecule for Toll-like receptor (TLR) signaling(*24*), we tested whether TLRs were required for the activity of CPL1. We found that deletion of Unc93b1, which is required for endosomal nucleic acid-sensing TLRs to function (*25*), did not impact CPL1-dependent arginase-1 induction (data not shown). However, we found abrogation of arginase-1 induction in response to CPL1 in the absence of the LPS receptor TLR4, while TLR2-deficiency had no effect (**Fig. 3H and 3I**). There are molecules that activate TLR4 signaling without directly binding the receptor such as palmitic acid (*26*). Notably, while TLR4-binding agonists direct rapid TLR4 endocytosis (*27*), indirect activators such as palmitic acid do not result in TLR4 internalization (*26*). To test the impact of CPL1 in TLR internalization, we stained BMDMs stimulated for 1hr with either LPS or rCPL1 and measured TLR4 surface levels by flow cytometry. We found that rCPL1 stimulation decreased surface TLR4 levels (**Fig. 3J**), though not to the same extent as LPS, consistent with a model in which CPL1 is a TLR4 ligand.

While our data did not find arginase-1 induction by LPS, we nonetheless addressed whether CPL1 phenotypes could be due to LPS contamination. First, we stimulated BMDMs with dilutions of LPS and tested arginase-1 upregulation. We found that LPS only induced robust arginase-1 at a high concentration (1ug/ml) (**Supp Fig. 4C**). As the high amount of LPS required for arginase-1 upregulation was at a concentration that induced robust caspase-11-dependent pyroptosis when given with cholera toxin B (CTB) (**Supp Fig. 4D**), we reasoned that if the effect of CPL1 were due to LPS contamination, we would observe pyroptosis when stimulating BMDMs with CPL1 + CTB(*28, 29*). However, we observed no pyroptosis induced by CPL1+CTB stimulation (**Supp Fig. 4E**). LPS is heat stable, whereas proteins can be denatured by heat. We observed that boiling of rCPL1 completely abrogated arginase-1 induction and potentiation by IL-4, again consistent with a role for CPL1 protein and not LPS (**Supp Fig. 4F**). Lastly, we tested whether expression of CPL1 in BMDMs could phenocopy the impact of the purified recombinant protein, as this would prevent the possibility of introducing microbial contaminants. We cloned CPL1 into a murine retroviral vector with an IL-2 signal peptide and transduced mouse bone marrow. After differentiation into BMDMs, we stimulated these cells with dilutions of IL-4. We found that transduction of CPL1 into macrophages potentiated the IL-4 induction of arginase-1 compared to control transductions (**Supp Fig. 4G**). Altogether, these data strongly support the conclusion that CPL1 activates TLR4 and this is not due to inadvertent endotoxin contamination.

While best known as a sensor of LPS on gram-negative bacteria, TLR4 plays a key role in driving allergic inflammation in response to house dust mite extract (HDM) and low dose LPS can also drive allergic inflammation in response to ovalbumin(*30, 31*). HDM is a complex mixture of proteins, but two dominant allergens are the protease Derp1 and the lipid-binding protein Derp2(*32, 33*). One mechanism by which Derp2 drives allergic responses is by acting as an MD2 mimic to activate TLR4(*34*). To test whether CPL1 could activate TLR4 via a similar mechanism, we reconstituted NF-kB-luciferase reporter expressing HEK293T cells with murine TLR4 alone or in combination with co-receptors CD14 and MD2 and then stimulated these cells with rCPL1 or LPS. While LPS only drove luciferase expression when TLR4 was co-expressed with CD14 and MD2, rCPL1 induced luciferase in cells expressing TLR4 only, suggesting that CPL1 can bypass a requirement for MD2 in TLR4 activation (**Fig. 3K**). In previous studies on Derp2, it was found that mutation of a key tyrosine residue flanked by two cysteines, which was similar to a conserved motif found on MD2, abrogated TLR4 activation, despite a lack of detectable sequence homology between the two proteins(*34*). We analyzed the CPL1 amino acid sequence and generated a recombinant protein with a mutation (Y160A) in an analogous motif (**Supp Fig. 4H**). Strikingly, we found that this mutation eliminated the ability of CPL1 to augment both IL-4-independent and -dependent arginase-1 expression (**Fig. 3L**). While it remains to be demonstrated whether CPL1 indeed activates TLR4 via an analogous mechanism to that of Derp2, this result provided an additional control for possible contaminants.

We next examined whether CPL1 contributes to type 2 inflammation during *in vivo* pulmonary infection. Deletion of IL-4 signaling in myeloid cells has been shown to be beneficial to the host during *Cryptococcus* infection, however, the lung myeloid compartment is highly heterogenous and the specific identity of the arginase1+ cells during fungal infection remains unclear(*35, 36*). We infected arginase-1-YFP reporter mice (YARG) with wild-type *C. neoformans* and processed lung tissue for flow cytometry on day 10. Analysis of the CD45+YARG+ cells revealed that interstitial macrophages (IMs) (CD45+CD64+MerTK+SiglecF-) are the major cell type that expresses Arg1 during *Cryptococcus* infection (**Fig. 4A and 4B**). To test the role of CPL1, we infected YARG mice intranasally with either wild type, *cpl1Δ*, or *qsp1Δ*, a strain lacking a secreted peptide previously shown to be important during *in vivo* infection(*37*). We found that *cpl1Δ* infection resulted in a striking reduction in YARG+ interstitial macrophages compared to both wild type and *qsp1Δ* infections (**Fig. 4C**). To determine whether this decrease in YARG+ cells was secondary to a global reduction in type 2 immunity we assessed other outputs of IL-4/IL-13 signaling such as eosinophilia, IgG1 class switching in germinal center B cells, and cytokine production by CD4+ T cells. Compared to wild type and *qsp1Δ, cpl1Δ* infection showed a modest reduction in lung eosinophils (**Supp Fig. 5A**), which is not surprising given that IL-4 stimulated macrophages produce eosinophil-recruiting chemokines such as CCL24 (*38*). On the other hand, we observed no difference between any fungal genotypes in the levels of IL-4-dependent IgG1 class switching (**Supp Fig. 5B and 5C**) or in CD4+ T cell cytokine production (**Supp Fig. 5D and 5E**). These data identify a crucial role for CPL1 in promoting local type 2 inflammation in the lung, although we cannot rule out effects on additional cell types beyond IMs.

**Figure 4.**
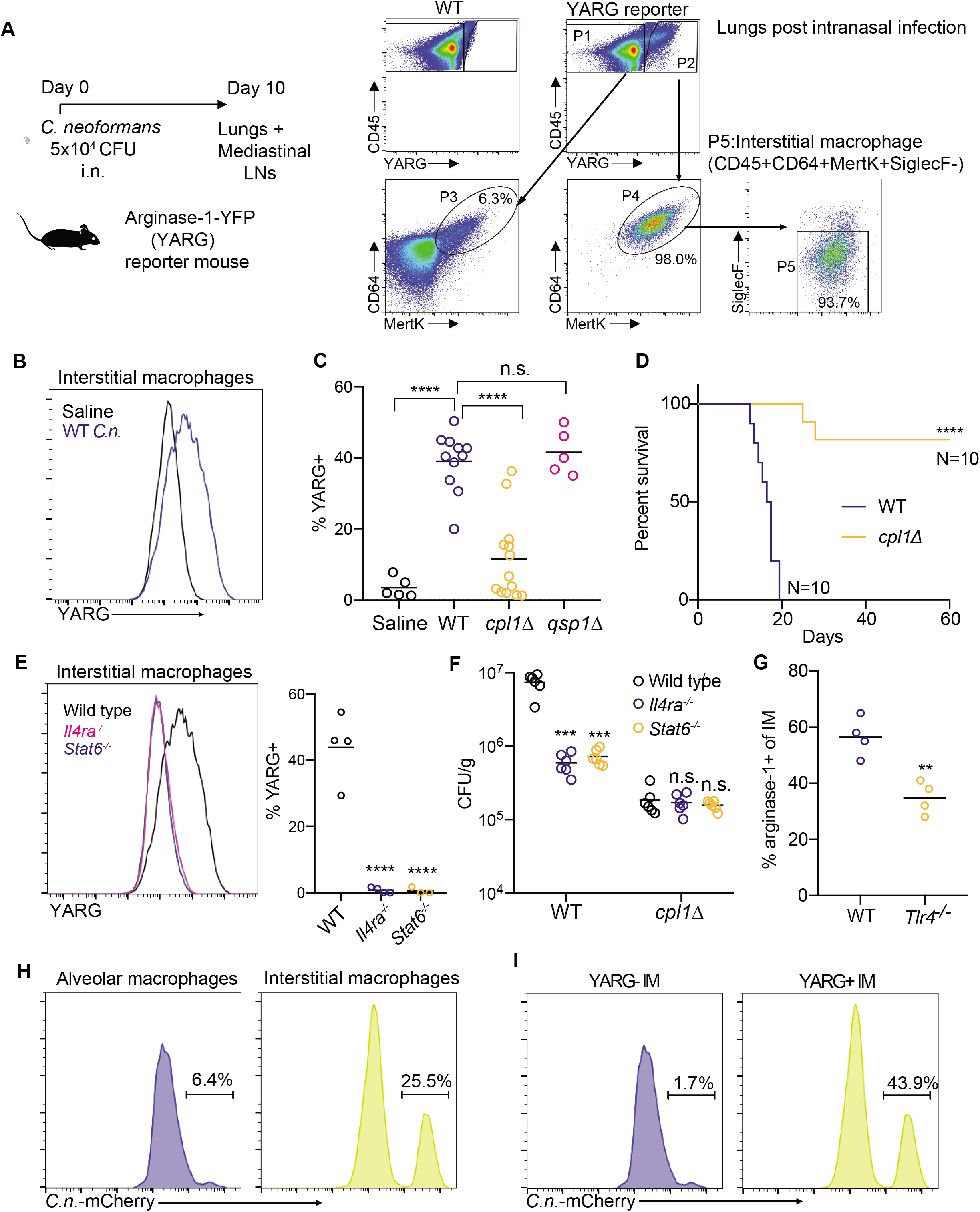
CPL1 promotes arginase-1 expression in pulmonary interstitial macrophages and is required for virulence. **(A)** FACS subgating on CD45+YARG+ lung cells from arginase-1-YFP (YARG) mice infected intranasally with 5×10^4^ CFU WT *C. neoformans* for 10 days. **(B)** Representative histogram of YARG expression in lung interstitial macrophages from mice injected intranasally with either saline or 5×10^4^ CFU WT *C.n*. for 10 days. **(C)** Quantification of YARG expression by FACS on interstitial macrophages from mice injected intranasally with either saline (N=5 mice), WT (N=11 mice), *cpl1Δ* (N=14 mice), or *qsp1Δ* (N=5 mice) Kn99a (5×10^4^ CFU) for 10 days; statistical significance determined by one-way ANOVA with Bonferroni test. **(D)** Kaplan-Meier survival curve analysis of mice infected with WT (N=10mice) or *cpl1Δ C.n*. (N=10 mice); ****p < 0.0001 by Mantel-Cox test. **(E)** Representative histogram (left) and quantification of YARG expression on WT, *Il4ra*^-/-^, or *Stat6*^-/-^ (all N=4 mice) infected for 10 days as in **(A)**. **(F)** CFUs from lungs of the indicated mouse genotypes (N=6 mice for each genotype) infected for 10 days with either WT or *cpl1Δ* strains; statistical significance determined by one-way ANOVA with Bonferroni test. **(G)** Arginase-1 FACS on lung IMs from WT or *Tlr4*^-/-^ mice infected as in **(A)**; statistical significance determined by unpaired T-test. **(H)** Representative FACS histograms of *C.neoformans*-mCherry expression in alveolar macrophages (left) or interstitial macrophages (right) after 10 days of infection. **(I)** Representative FACS histograms of *C.neoformans*-mCherry expression in interstitial macrophages from YARG mice gated on YARG-negative IMs (left) or YARG-positive IMs (right). *p < 0.05; **p < 0.01; ***p < 0.001; ****p < 0.0001

A defect in type 2 cytokine signaling increases survival of mice infected with *C. neoformans* (*35*). Since we found that mice infected with *cpl1*Δ *C. neoformans* exhibited decreased type 2 inflammation, we asked whether CPL1 impacted mouse survival. Strikingly, we found that nearly all mice survived infection by a strain lacking CPL1 (**Fig. 4D**). Complementation of the mutant with the wild-type gene rescues the *in vivo* YARG and eosinophil defects as well as the virulence defect (**Supp Fig. 6A, 6B, 6C**). We confirmed that YARG induction in response to *C. neoformans* infection was indeed dependent on type 2 cytokine signaling using both IL-4Rα- and STAT6-deficient mice (**Fig. 4E**).

As CPL1 may have contributions to virulence *in vivo* beyond augmentation of type 2 inflammation, we tested whether the *cpl1*Δ mutant displayed phenotypes in mice lacking factors critical for type 2 responses. We infected wild type, *Il4ra*^-/-^, and *Stat6*^-/-^ mice with either wild type or *cpl1Δ C. neoformans* and then measured pulmonary CFUs on day 10. Consistent with previous reports, we found that IL-4Rα- and STAT6-deficient mice showed lower pulmonary fungal burden than wild type mice (**Fig. 4F**). However, while *cpl1Δ* infections showed decreased CFUs relative to the control strain in wild type mice, there was no further decrease in IL-4Rα- and STAT6-deficient hosts (**Fig. 4F**). This result suggests that induction of type 2 inflammation is required for CPL1 to impact infectivity. We reasoned that simultaneous infection of mice with wild type *C. neoformans* might rescue *cpl1Δ* CFUs via the induction of type 2 immunity in *trans*. We performed mixed intranasal infections [WT vs. WT(KanR), *cpl1Δ* vs. *cpl1Δ* (KanR), and WT(KanR) vs. *cpl1Δ*] using congenic G418-resistant mixing partners and found that co-infection with wild type partially restored the *cpl1Δ* pulmonary CFUs (**Supp Fig. 6D**). While there are likely additional mechanisms by which CPL1 promotes *C. neoformans* pathogenesis, these data indicate that promotion of type 2 immunity is a critical function of this virulence factor.

Since BMDM polarization by CPL1 requires TLR4, we tested whether TLR4 contributed to type 2 inflammation in response to pulmonary infection. We found that *Tlr4*^-/-^ mice infected with *C. neoformans* displayed reduced arginase-1 expression in IMs as well as reduced pulmonary eosinophils (**Fig. 4G and Supp Fig. 6E**). To investigate the role of CPL1 in animals but in the absence of pathogen-host dynamics, we tested whether rCPL1 protein alone could drive type 2 inflammation. We challenged wild type and *Tlr4*^-/-^ mice intranasally with a series of CPL1 doses and then assessed pulmonary eosinophilia on day 21. We found that intranasal treatment of mice with rCPL1 (but not bovine serum albumin) was sufficient to induce pulmonary eosinophilia; this induction was fully dependent on TLR4 (**Supp Fig. 6F**).

Alternatively activated macrophages can be exploited as a replicative niche by pathogens due to their reduced ability to kill microbes (*39*). To begin to test this hypothesis we performed flow cytometric analysis of pulmonary tissue from YARG mice infected intranasally with mCherry-expressing *C. neoformans*. We observed high mCherry+ signal associated with interstitial macrophages compared to alveolar macrophages (**Fig. 4H**). Sub-gating interstitial macrophages into either YARG- or YARG+ populations further showed a striking enrichment of mCherry+ *C. neoformans* in the YARG+ cells (**Fig. 4I**). These data demonstrate a selective association of *C. neoformans* with arginase-expressing IMs *in vivo*.

Using forward genetics, we identified CPL1 as a secreted effector of *C. neoformans* that promotes fungal virulence by enhancing type 2 inflammation. Our work supports a model whereby CPL1 activates TLR4 signaling to drive phosphorylation of STAT3 in macrophages, which both promotes the initial induction of arginase-1 but also amplifies macrophage sensitivity to IL-4 signaling (**Supp Fig. 6G**). *In vivo*, CPL1 is required for virulence and, strikingly, promotes the induction of arginase-1 by interstitial macrophages. *C. neoformans* also physically associates with polarized IMs during infection, consistent with direct macrophage reprogramming *in vivo* by CPL1. While detailed biochemical and structural studies will be required understand the precise mechanism of TLR4 activation by CPL1, it is nonetheless tempting to speculate that CPL1 might deliver a fungal-derived ligand to this receptor analogous to MD-2-LPS. The immunomodulatory functions of CPL1 revealed here demonstrate that a human pathogenic fungus produces an effector molecule that interfaces directly with mammalian immune cells. We anticipate that ongoing advances in fungal pathogen genetics will lead to the identification of additional secreted effectors and host targets. In turn, such studies will begin to reveal the constellation of mechanisms that underpin the host immune reprograming events that drive the virulence of invasive fungal pathogens, providing new opportunities to combat these highly lethal infections.

## Acknowledgements

We thank Prof. Gregory Barton for kind provision of Tlr2-, Tlr4-, and Tlr2×4-knockout mice, and Prof. Roberto Ricardo-Gonzalez for provision of *Il4ra*^-/-^ and *Stat6*^-/-^ mice. We thank Prof. Seemay Chou for helpful advice on protein purification. We thank Prof. Jason Cyster and Prof. Emily Goldberg for critically reading the manuscript, and we thank them as well as Drs. Sandra Catania, Michael Boucher for discussions and advice.

## Author Contributions

Conceptualization: EVD, HDM

Methodology: EVD, AR, HDM

Investigation: EVD, SL

Visualization: EVD

## Supplementary materials

### Materials and methods

#### Mice

Wild type C57BL/6J mice were purchased from Jackson Labs and bred in house. *MyD88*^-/-^ and *Stat3^flox/flox^* mice were purchased from Jackson Labs. Arginase-1-YFP (YARG) reporter mice, *IL-4ra*^-/-^ and *State6*^-/-^ mice were a gift from Richard Locksley (UCSF). BoyJ (B6.SJL-*Ptprc^a^Pepc^b^*/BoyJ) mice were a gift from Jason Cyster (UCSF). Animals were housed in a specific-pathogen free environment in the Laboratory Animal Research Center at UCSF. All experiments conformed to ethics and guidelines approved by the UCSF Institutional and Animal Care and Use Committee.

#### Yeast manipulations

Yeast genetic manipulations were performed as previously described {Chun:2010er}. Insertion of genes was obtained through homologous recombination by transforming 10ug of digested plasmid.

#### Intranasal infections

Individual colonies from *Cryptococcus* plates were cultured overnight in 10mL of YPAD (30C). The next day, yeast were counted on a hemocytometer and diluted to 1×10^6^ cells per mL in saline. Mice were anesthetized with intraperitoneal ketamine/dexmedetomidine and then hung by their front teeth using surgical thread. 50uL of yeast (5×10^4^ CFU) were then pipetted onto the nasal flares and taken up by aspiration. For survival curves, the mice were weighed and assessed for clinical endpoints every 2 days for the first week post-infection, and then every day starting after week 1.

#### Intranasal injection of CPL1

Mice were anesthetized with intraperitoneal ketamine/dexmedetomidine and then hung by their front teeth using surgical thread for intranasal injection. Mice were injected with 100ng of either bovine serum albumin (Sigma) or recombinant CPL1 on day 0, 1, and 2 and then injected with 25ng of either protein on days 14, 15, 17, 18, 19 and then mice were euthanized for analysis on day 21.

#### Generation of bone marrow-derived macrophages

On day 0, mice were euthanized and femurs/tibias were harvested into RPMI (2% FCS). The ends of the bones were clipped with scissors, and then bone marrow was flushed using a 25^1/2^ gauge syringe. The bone marrow was then suspended in complete DMEM (10% FCS, HEPES, glutamine, P/S) and 10% m-CSF media derived from 3T3-mCSF cells. Bone marrow suspensions were plated in non-tissue culture-treated 10cm petri dishes. On day 4, 3mL of additional 10% m-CSF media was added to each plate. The adherent macrophages were then harvested on day 6 and either plated out for experiments or frozen.

#### Retroviral transduction of BMDMs

On day 0, bone marrow was harvested and cultured in 10% m-CSF media as above and murine retroviral MSCV plasmid encoding CPL1 or iCre was transfected into platE cells using Lipofectamine 2000 (Thermo Fisher Scientific). On day 2, viral supernatants were harvested and filtered through a 0.45micron syringe filter. Non-adherent bone marrow cells were harvested and spun down in 6-well plates. The supernatant was aspirated, and 2mL of retrovirus was added to each well along with 10ug/ml polybrene (Sigma). The plates were then spun for 2hrs at room temperature at 2400rpm with no brake. After spinning, the retroviral supernatants were aspirated and replaced with 10% m-CSF media. On day 3, another round of identical spinfection was performed. Cells were then harvested for experiments on day 6.

#### ELISA

High-binding half area 96-well plates (Corning) were coated with 25uL of unconjugated anti-TNF antibody (Invitrogen; clone 1F3F3D4) at a concentration of 2ug/ml in PBS and incubated overnight at 4°C. Plates were then washed 6 times with PBST (1X PBS + 0.05% Tween-20) and blocked for 1hr at room temperature with 120uL of 1X PBS + 5% FCS. Next, the blocking solution was removed and 25uL of indicated macrophages supernatants plus a standard curve using recombinant murine TNF (Peprotech) in 1xPBS + 5% FCS were added to the plates and incubated for an hour at room temperature. Plates were washed as above, and then 25uL of biotinylated anti-TNF (Invitrogen; clone MP6-XT3) was added a concentration of 1ug/ml in PBS + 5% FCS for 1hr at room temperature. Plates were then washed, and 25uL of streptavidin-HRP (Jackson ImmunoResearch) was added at a concentration of 2.5ug/ml in 1xPBS + 5% FCS for 1hr at room temperature. Plates were then washed, assay was developed using a 50uL of Substrate reagent (R&D Systems), and absorbance was read at 450nm.

#### Flow cytometry

Cells were stained with Abs to CD11b (M1/70), SiglecF (S17007L), Arginase-1 (A1exF5), iNOS (CXNFT), Mertk (DS5MMER), CD64 (X54-5/7.1), CD90.2 (53-2.1), CD45.2 (104), CD45.1 (A20), B220 (RA3-6B2), CD38 (NIMR5), IgD (11-26c.2a), CD4 (GK1.5), CD95 (Jo2), GL7 (GL7), IgG1 (RMG1-1), IgA (C10-1), IgM (11/41), IL-4 (11B11), IL-17A (TC11-18H10.1), IFNg (XMG1.2), TCRβ (H57-597), CD44 (IM7), TLR4 (SA15-21), IL-4Rα (I015F8), GXM (18B7) (from Biolegend, BD Biosciences or eBiosciences). To detect intracellular arginase-1 or iNOS, cells were treated with BD Cytofix Buffer and Perm/Wash reagent (BD Biosciences) and then stained with anti-arginase-1 or anti-iNOS in Perm/Wash buffer. For flow cytometry on lung samples, mice were infected or challenged as indicated and then lungs were dissected and minced using scissors. The minced lung tissue was then incubated for 30min at 37C in digestion media (RPMI, 2% FCS, 0.125mg/ml Collegenase II (Thermo Fisher Scientific), 0.2mg/ml DNaseI (Millipore)). The digested lung tissue was then mashed through a 100micron strainer (Fisher Scientific) and washed with RPMI + 2% FCS + 5mM EDTA. Red blood cells were then lysed for 5min on ice using RBC Lysis Buffer (Biolegend). Cells were resuspended in FACS buffer (1XPBS, 2% FCS, 1mM EDTA) for staining. For analysis of mediastinal lymph nodes, the LNs were dissected from mice and then mashed through a 100micron strainer.

#### Expression of recombinant protein in *Pichia pastoris*

CPL1 was PCR amplified from Kn99a genomic DNA with a GSGS-linker-6xHis tag encoded in the 3’ primer. This PCR product was then cloned into SnaBI-digested pPIC9K (Thermo Fisher Scientific) using a Gibson assembly kit (New England Biosciences). The assembled vector was transformed in to DH5α *E. coli* and plated onto LB + ampicillin plates. Colonies were screened for the correct inserts by sanger sequencing (Quintara Bio). For *P. pastoris* transformation, a 50mL culture of GS115 was inoculated overnight at 30°C at 200rpm. The next morning, the cultures were diluted into 500mL YPAD and incubated (30°C, 200rpm) until the culture reached OD600=2.0. Then cultures were then spun down and washed twice with ice cold 1M sorbitol. The cells were then resuspended in 2mL of ice cold sorbitol and then electroporated with 5ug of *SacI*-digested pPIC9K-CPL1-6xHis. Electroporated yeast were then selected for successful integrations on HIS-plates.

#### Purification of recombinant CPL1

Single colonies were inoculated in 100mL of BMGY (1% yeast extract, 2% peptone, 100mM potassium phosphate pH 6.0, 1.34% YNB, 4×10^-5^% biotin, 1% glycerol) overnight at 30C, 200rpm. The next day, yeast were pelleted, washed with ddH_2_O and then resuspended to OD_600_=1.0 in BMMY (1% yeast extract, 2% peptone, 100mM potassium phosphate pH 6.0, 1.34% YNB, 4×10^-5^% biotin, 1% methanol) for induction. Supernatants from BMMY induction cultures (24hrs at 30°C, 300rpm) were concentrated using 10kD-cutoff Centricon Plus −70 concentrators (EMD Millipore). The concentrated supernatants were then dialyzed for 24hrs (10mM Phosphate buffer pH=7.4, 500mM NaCl, 10% glycerol) using 30mL 10kD pore Slide-a-Lyzer Dialysis cassettes (Thermo Fisher Scientific). The dialyzed supernatants were then run over a 5mL HisTrap HP column using an ĀKTA pure fast protein chromatography system (Cytiva). The column was equilibrated with 25mL of equilibration buffer (10mM Phosphate buffer pH=7.4, 500mM NaCl, 10% glycerol, 20mM imidazole) and then the dialyzed supernatants were injected onto the column. The column was then washed with 25mL of equilibration buffer. The bound proteins were then eluted in 15mL of elution buffer (10mM Phosphate buffer pH=7.4, 500mM NaCl, 10% glycerol, 500mM imidazole). The eluted proteins were concentrated using 10kD cutoff Amicon Ultra-15 Centrifugal Filter units (EMD Millipore) and then further purified on a HiLoad 16/600 Superose 6pg preparative size exclusion chromatography column (SEC) (GE Healthcare). The SEC column was first equilibrated with 128mL of equilibration buffer (10mM Phosphate buffer pH=7.4, 125mM NaCl, 10% glycerol) and then the loaded 1mL sample was injected, and fractions were collected over a 128mL elution volume. The eluted fractions were run on a 4-12% SDS-PAGE (Fisher Scientific) gel and analyzed by silver stain and western blot for purity. Fractions containing pure CPL1-6xHis were then concentrated to 2mg/ml using 10kD cutoff Amicon Ultra-4 Centrifugal Filter units (EMD Millipore).

#### RNA-seq

2×10^6^ BMDMs were seeded in 6-well plates and stimulated with the indicated conditions. RNA was then extracted from the macrophages using a RNeasy Midi Kit (Qiagen). Sequencing libraries were prepared using 500ng of purified RNA using a QuantSeq 3’end mRNA-Seq Library Prep Kit (Lexogen). Library quality and quantify was determined using a High Sensitivity DNA Bioanalyzer chip (Agilent). The RNA-seq libraries were then sequenced using 50bp single end reads on a HiSeq4000 (Illumina).

#### RNA-seq analysis

Read counts were determined using HTSeq by counting the number of reads aligned by STAR for each mouse transcript. We then used DEseq2 to determine differentially expressed genes between different treatment conditions.

#### Screening the *Cryptococcus* knockout collection

Individual Cryptococcus knockout strains were spotted onto YPAD+NAT omni-trays in a 96-well pattern (Thermo Fisher Scientific) from frozen −80C stocks. Each mutant was then inoculated into 100uL of YPAD in 96-well round bottom plates and incubated overnight at 30C on a shaker. The OD_600_ was then determined for each well using a plate reader, and yeast were diluted into complete DMEM at a concentration of 1×10^7^ cells/ml. 100ul of each mutant was then added to BMDMs (1×10^5^ cells) seeded in 96-well plates. Infections were left for 24hrs, and then the cells were surface stained for CD11b to distinguish macrophages from yeast, and then intracellular argainse-1 staining was performed as described above. The screen was performed over several rounds of experiments, with an average of 5 knockout plates screened per experiment. For each experiment, a 96-well plate of entirely wild type Cryptococcus was inoculated and added to macrophages. Z-scores for the mutants in each experiment were calculated using the mean and standard deviation for arginase-1 induction by the wild type plate.

#### Western blotting

BMDMs were stimulated with the indicated conditions and then supernatants were aspirated and cells were washed with ice cold 1XPBS. The cells were then placed on ice and lysed for 20min in RIPA buffer (50mM Tris-HCl pH=7.4, 150mM NaCl, 1% NP-40, 0.5% deoxycholate, 0.1% SDS) plus protease/phosphatase inhibitor cocktail (Thermo Scientific). The lysate was then transferred to 1.5mL Eppendorf tubes and centrifuged for 20min at 14,000rpm at 4C. Novex NuPAGE LDS samples buffer (Thermo Fisher Scientific) was then added and samples were run on 4-12% NuPAGE Bis-Tris protein gels (Thermo Fisher Scientific) at 190V. Gels were then transferred onto nitrocellulose membranes at 35V for 90min. Membranes were blocked for 1hr in 5%(w/v) milk in TBST. Primary antibodies were then added at 1:1000 in TBST + 5% mil overnight at 4C. Antibodies were used against Stat6 (D3H4), pStat6-Tyr641 (D8S9Y), Stat3 (79D7), pStat3-Tyr705 (D3A7) (all purchased from Cell Signaling Technologies). Membranes were then washing 3×5minutes using TBST and secondary anti-rabbit horseradish (Bio-Rad) peroxidase was added at 1:10,000 for 1hr at room temperature. Membranes were washed as above and then developed for 5min using a SuperSignal West Pico Plus Chemiluminescent Substrate kit (Fisher Scientific).

#### LDH release assay

1×10^5^ BMDMs were seeded in 96-well plates and then stimulated with the indicated conditions in DMEM without phenol red (Corning). Supernatants were then harvested and LDH release was determined using a LDH Cytotoxicity Assay kit (Fisher Scientific) in 96-well plates. The assay was then measured at 490nm absorbance. The percent LDH release was determined by comparing to positive control samples that were macrophage RIPA buffer lysates.

#### Total Nitric oxide assay

1×10^5^ BMDMs were seeded in 96-well plates and then stimulated with the indicated conditions. Supernatants were then assayed for nitric oxide in 96-well plates using a colorimetric Total Nitric Oxide Assay Kit (Thermo Fisher Scientific). Assays were read on a plate reader at an absorbance of 540nm.

#### India ink capsule staining

*C. neoformans* single colonies were picked from YPAD plates and grown overnight in YPAD at 30°C with shaking. The next day, cultures were diluted 1:100 in capsule induction media (10% Sabouraud, 50mM HEPES pH=7.9) and incubated overnight at 30°C with shaking. Yeast cells were then fixed with 2% paraformaldehyde for 15min at room temperature, and washed twice with 1XPBS. Cells were resuspended in 100uL of 1XPBS and then diluted 1:1 with India ink. Cells were then placed on slides with coverslips and imaged with a 40X objective lens using brightfield microscopy (Leica).

#### RT-qPCR

*C. neoformans* was cultured in the indicted media and temperatures until reaching OD_600_=1.0. Yeast were then pelleted, resuspended in 1mL Trizol (Thermo Fisher Scientific), and lysed using a bead beater (2 cycles x 90 seconds). 200uL of chloroform was then added and the suspension was vortexed until homogenous and centrifuged at 12,500xg at 4°C. The aqueous phase was then collected and RNA was further extracted using an RNA Extraction Kit (Zymo Research). Reverse transcription was then performed on 5ug of RNA using a SuperScript III™ Reverse Transcription Kit (Thermo Fisher Scientific). Quantitative PCR was then ran with PowerUP SYBR Green Master Mix (Life Technologies) to determine the expression of *CPL1* (Fwd: 5’-CTCGCAGACTGGTTCAAGGT-3’; Rev: 5’-GCGCAATCTTGGCCAGAAC-3’) relative to *ACT1* (Fwd: 5’-CCACCCACTGCCCAAGTAAA-3’; Rev: 5’-GTCGAGGGCGACCAACAATA-3’).

---

**Supplementary Figure 1.**
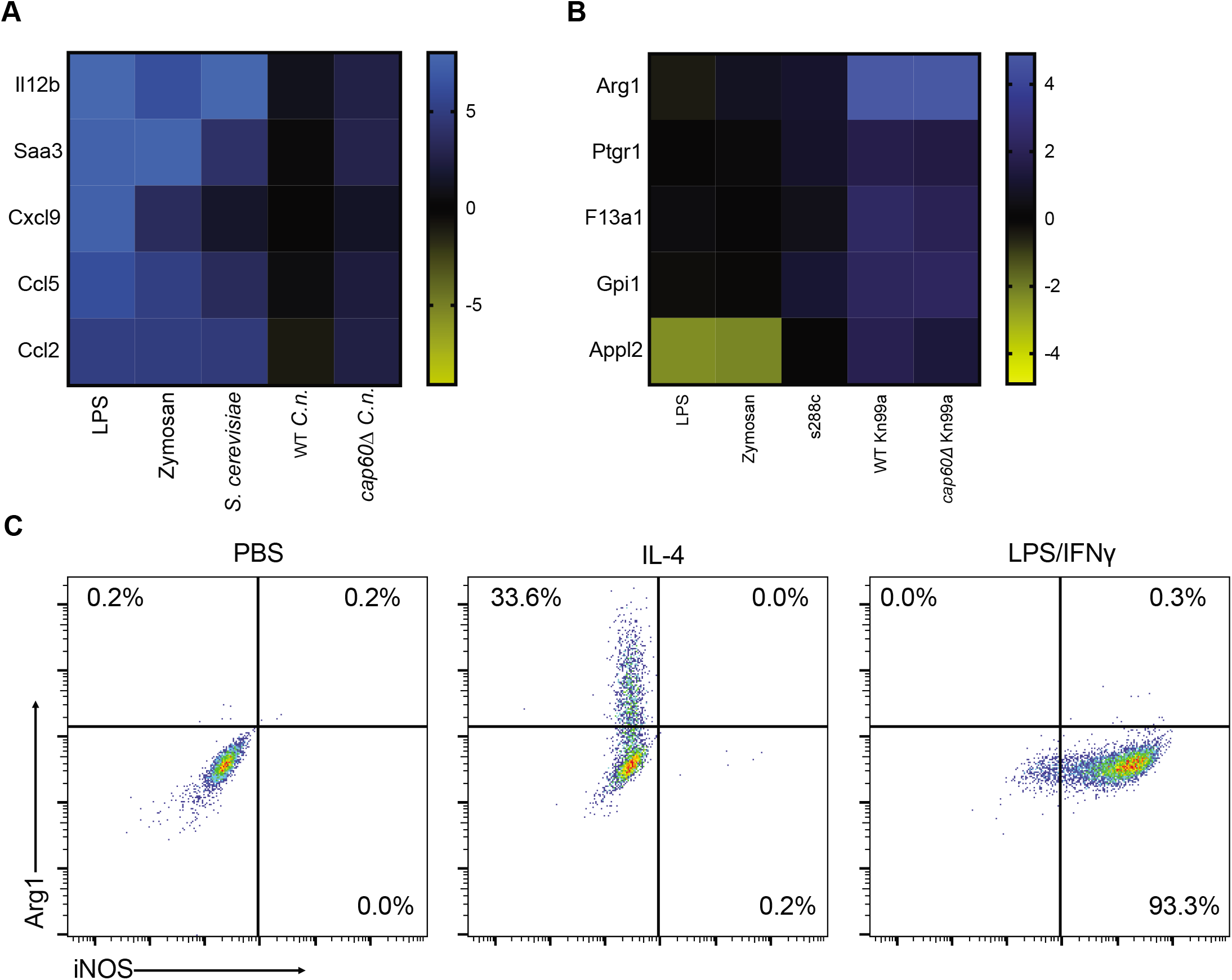
**(A)** RNA-seq heatmap depicting log_2_ fold changes of the indicated pro-inflammatory genes in BMDMs following 6hrs of stimulation. **(B)** RNA-seq heatmap depicting log_2_ fold changes of the indicated M2/tolerized genes in BMDMs following 6hrs of stimulation. **(C)** Representative FACS plots of Arg1 and iNOS expression in BMDMs following 24hrs of stimulation with PBS, IL-4 (40ng/ml), or LPS (100ng/ml) and IFNγ (50ng/ml).

**Supplementary Figure 2.**
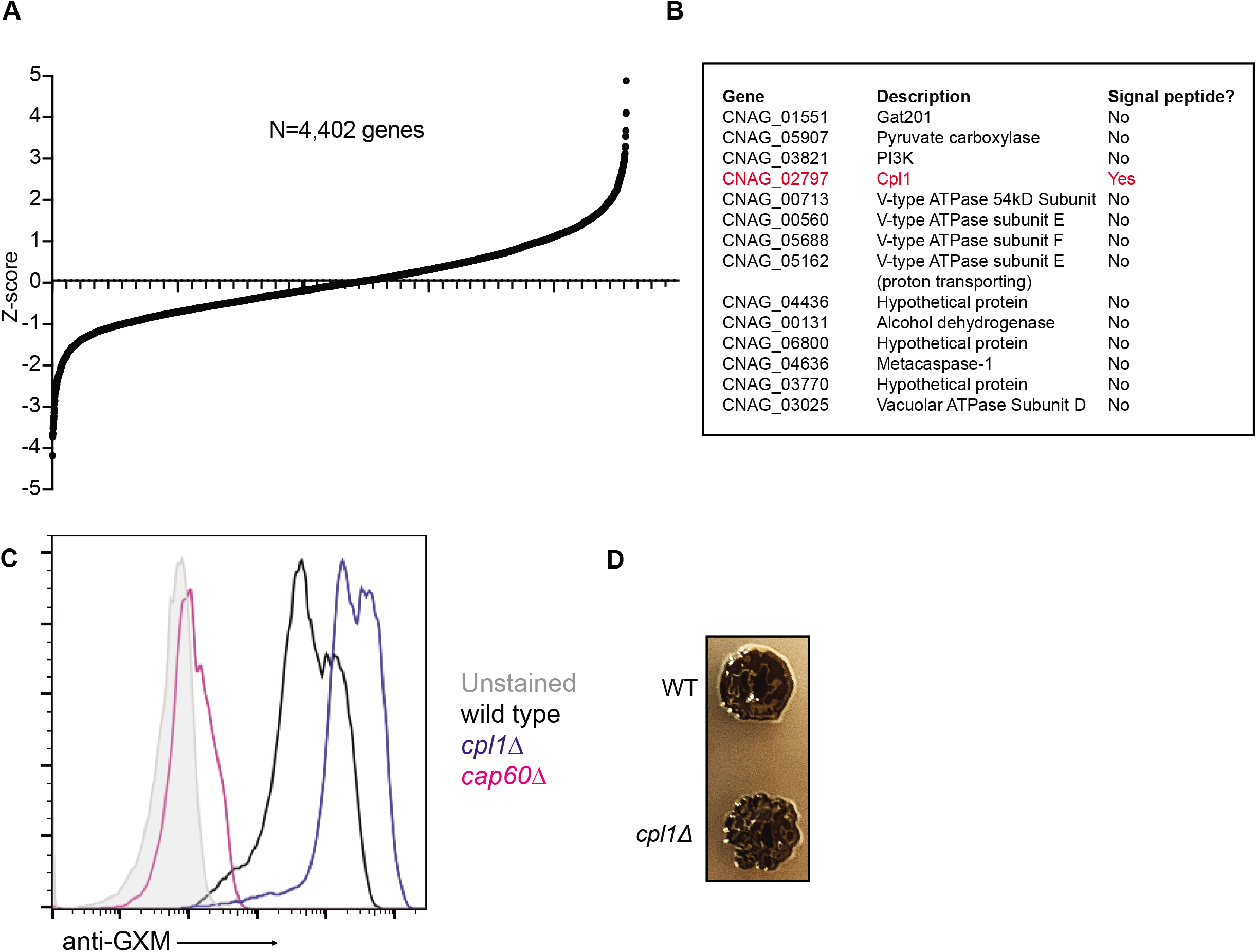
**(A)** Ranked Z-scores of hits from forward genetic screen for *C. neoformans* Arg1 induction. **(B)** List of validated screen hits and gene descriptions. **(C)** Representative FACS histograms of GXM staining on the indicated C. neoformans strains cultured overnight in 10% Sabouraud media. **(D)** Melanin production in WT or *cpl1Δ* strains grown at 30°C on L-DOPA plates.

**Supplementary Figure 3.**
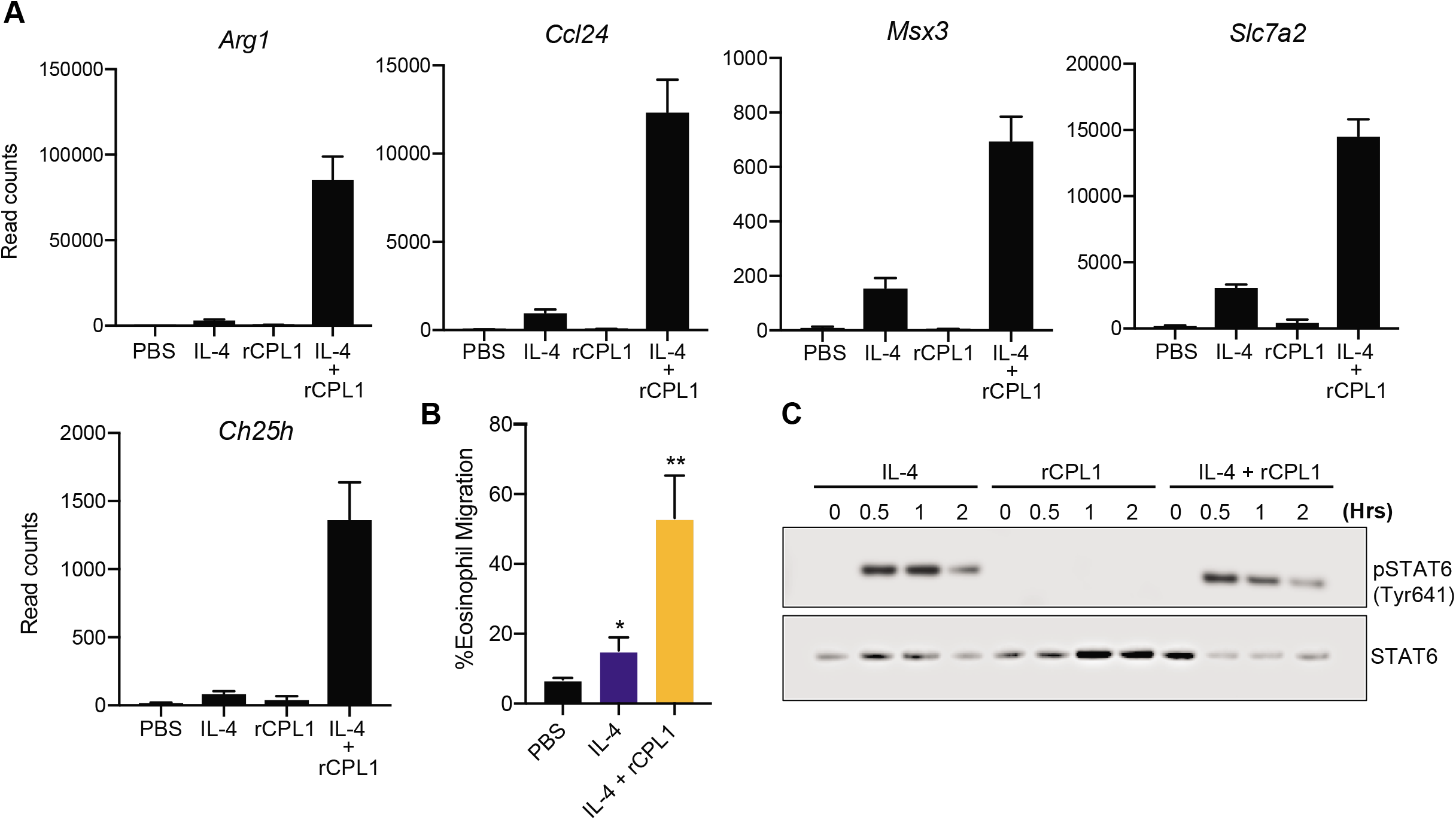
**(A)** RNA-seq read counts of the indicated genes in BMDMs stimulated for 24hrs with either PBS, IL-4 (10ng/ml), rCPL1 (111nM), or IL-4 + rCPL1. **(B)** Transwell migration assay on splenic eosinophils towards supernatants from BMDMs stimulated as in **(A)**. **(C)** Western blot for pStat6 or total Stat6 on BMDMs stimulated for the indicated times with either IL-4 (10ng/ml) alone, rCPL1 (111nM) alone, or IL-4 + rCPL1.

**Supplementary Figure 4.**
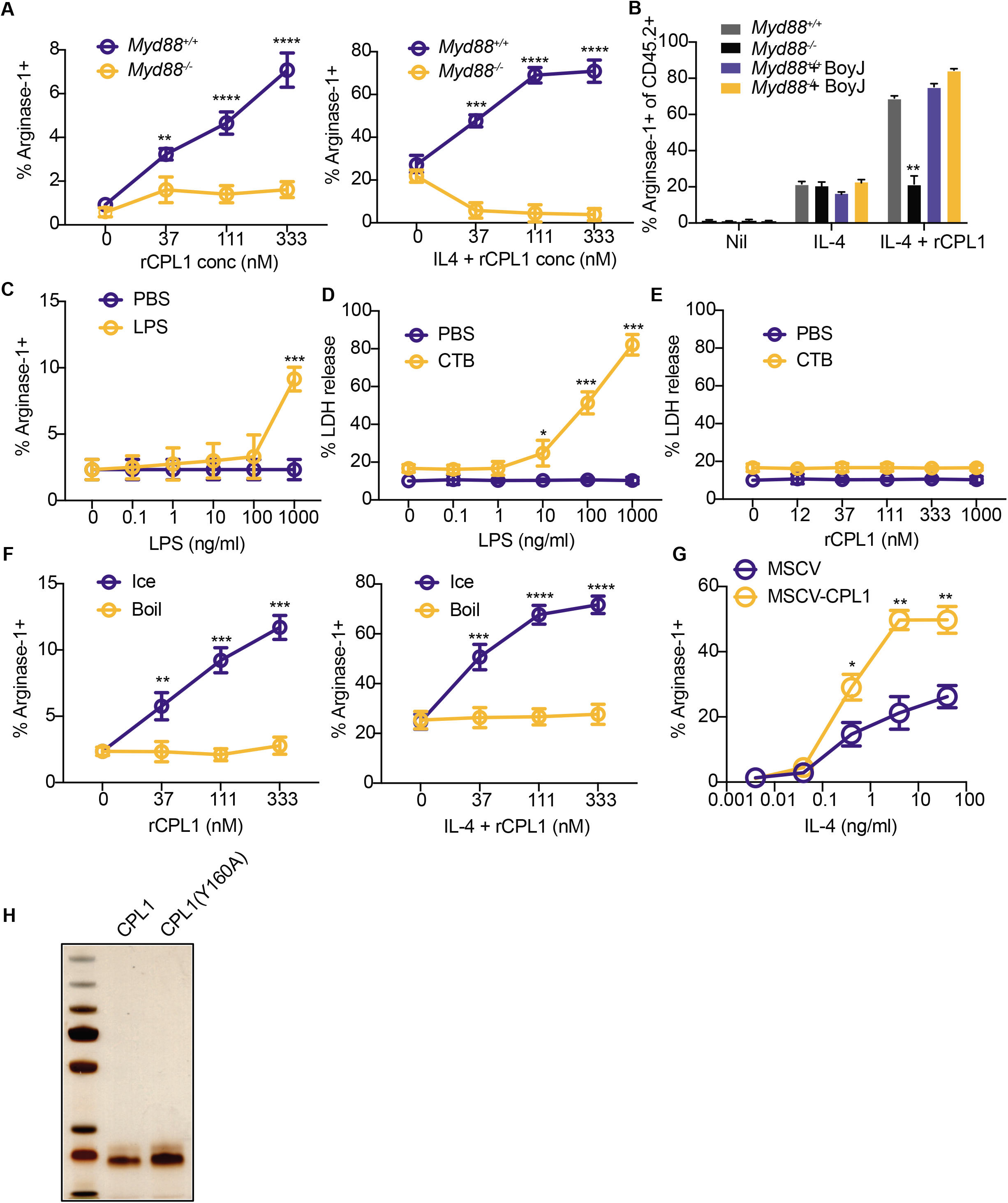
**(A)** Arginase-1 FACS in *Myd88*^+/+^ or *Myd88*^-/-^ BMDMs stimulated for 24hrs with the indicated concentrations of rCPL1 alone (left) or in combination with IL-4 (10ng/ml). **(B)** Arginase-1 FACS gated on CD45.2+ BMDMs from the indicated genotypes co-cultured with a 50:50 mix of CD45.1 BoyJ BMDMs and stimulated for 24hrs with IL-4 (10ng/ml) or IL-4 + rCPL1 (111nM). **(C)** Arginase-1 FACS on BMDMs stimulated for 24hrs with the indicated concentrations of LPS. **(D)** Measurement of pyroptosis by LDH release assay on BMDMs stimulated with the indicated concentrations of LPS alone or with 10ug/ml cholera toxin B (CTB). **(E)** Measurement of pyroptosis by LDH release assay on BMDMs stimulated with the indicated concentrations of rCPL1 alone or with 10ug/ml CTB. **(F)** Arginase-1 FACS on BMDMs stimulated with rCPL1 that was either kept on ice or boiled at 100°C for 15min. Cells were stimulated with either rCPL1 alone (left) or in combination with IL-4 (right). **(G)** Arginase-1 FACS on BMDMs transduced with MSCV-empty or MSCV-CPL1 retrovirus and stimulated for 24hrs with the indicated concentrations of IL-4. **(H)** Silver stain on SDS-PAGE gel of rCPL1-6xHis or rCPL1(Y160A)-6xHis purified from *P. pastoris*. *p < 0.05; **p < 0.01; ***p < 0.001; ****p < 0.0001 by one-way ANOVA with Bonferroni test.

**Supplementary Figure 5.**
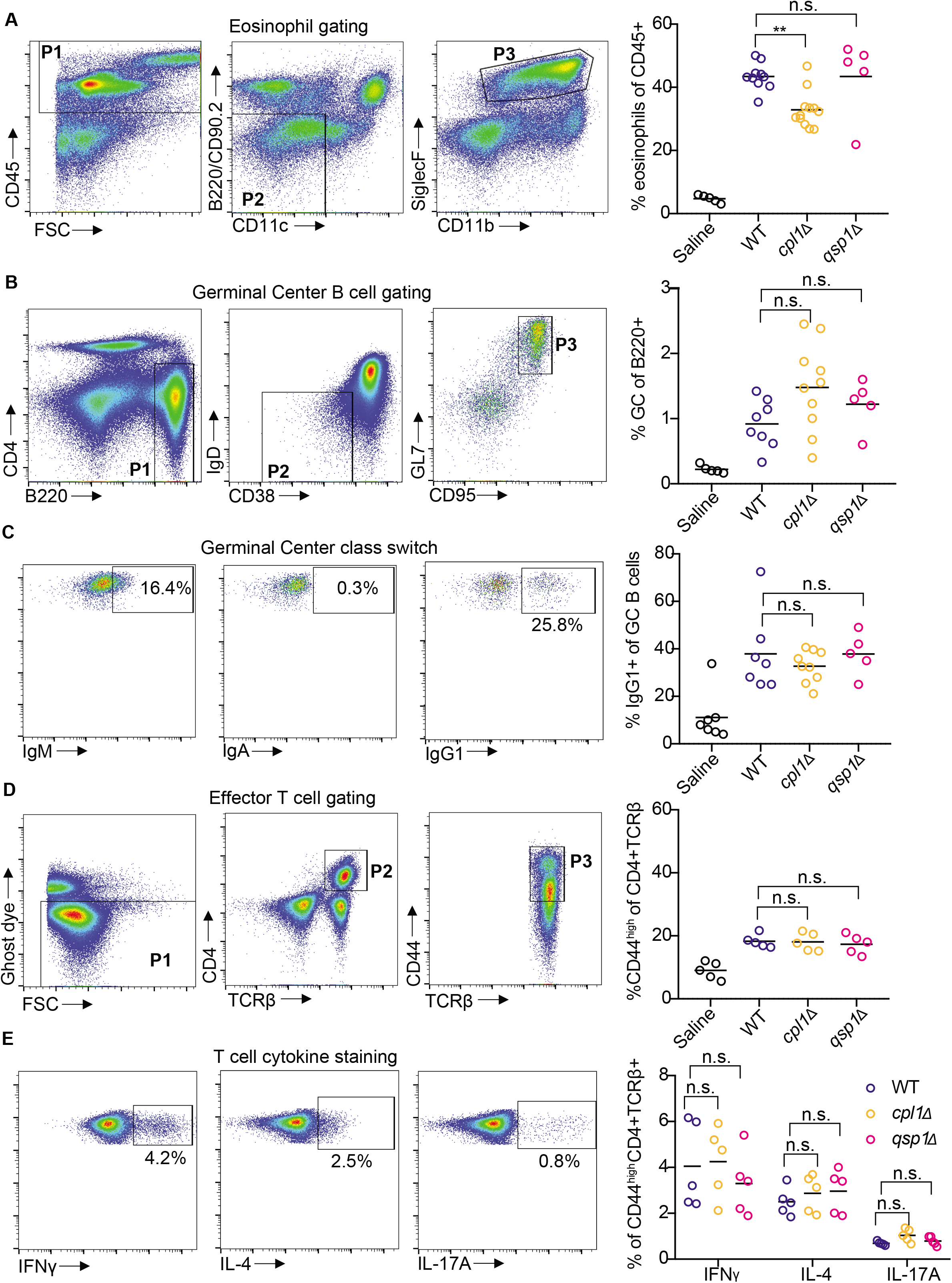
**(A)** Representative FACS gating and quantification of lung eosinophils after 10 days of intranasal infection with the indicated *C.n*. strains. **(B)** Representative FACS gating and quantification of mediastinal lymph node GC B cells. **(C)** Representative FACS gating and quantification of GC B cell antibody isotype. **(D)** Representative FACS gating and quantification of effector CD4+ T cells. **(E)** Representative FACS gating and quantification of cytokine production from effector CD4+ T cells after 4hrs of stimulation with PMA, Ionomycin, and GolgiSTOP. **p < 0.01 by one-way ANOVA with Bonferroni test.

**Supplementary Figure 6.**
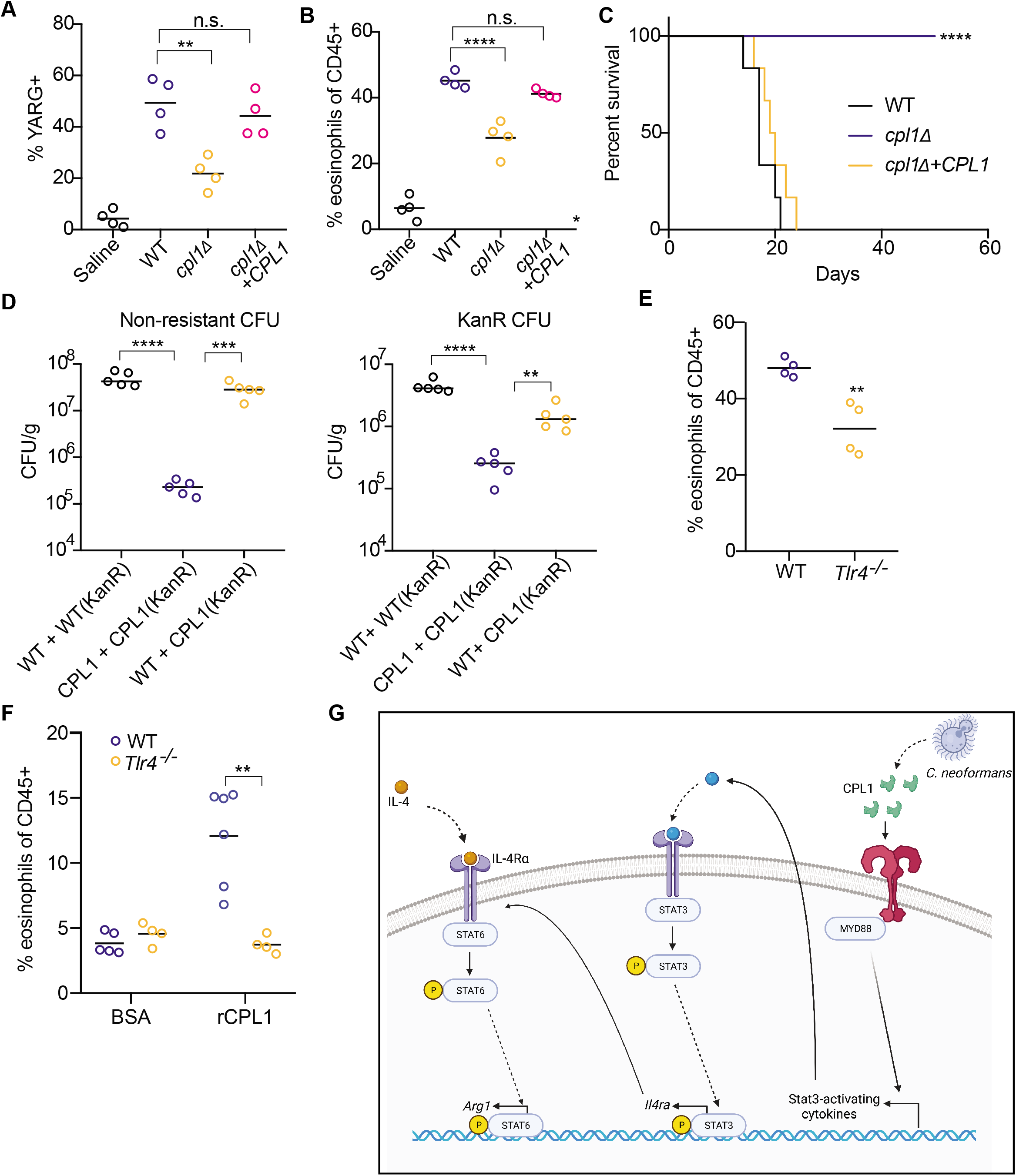
**(A)** Quantification of YARG expression by FACS in lung interstitial macrophages in mice infected for 10 days with 5×10^4^ CFU of the indicated strains. **(B)** Quantification of eosinophils in mice infected for 10 days with 5×10^4^ CFU of the indicated strains. **(C)** Kaplan-Meier survival curve analysis of mice infected with WT, *cpl1Δ*, or *cpl1Δ+CPL1 C.n*. (N=6 mice per group); ****p < 0.0001 by Mantel-Cox test. **(D)** Lung CFUs on G418-non-resistant (left) or -resistant (right) colonies from mice infected for 10 days with a 50:50 mix of the indicated strains. **(E)** Quantification of lung eosinophils in WT or *Tlr4*^-/-^ mice infected for 10 days with 5×10^4^ CFU *C. neoformans*. **(F)** FACS quantification of lung eosinophils in WT (N=6 mice) or *Tlr4*^-/-^ (N=4 mice) mice sensitized intranasally with rCPL1. **(G)** Model of how secreted CPL1 modulates the macrophage inflammatory state (created using Biorender.com). p < 0.05; **p < 0.01; ***p < 0.001; ****p < 0.0001 by one-way ANOVA with Bonferroni test.

